# Spatial architecture of CD8^+^ T cells and DC subsets is critical for the response to immune checkpoint inhibitors in melanoma

**DOI:** 10.1101/2024.02.06.579128

**Authors:** Elisa Gobbini, Margaux Hubert, Anne-Claire Doffin, Anais Eberhardt, Leo Hermet, Danlin Li, Pierre Duplouye, Sarah Barrin, Justine Berthet, Valentin Benboubker, Maxime Grimont, Candice Sakref, Jimmy Perrot, Garance Tondeur, Olivier Harou, Jonathan Lopez, Bertrand Dubois, Stephane Dalle, Christophe Caux, Julie Caramel, Jenny Valladeau-Guilemond

**Author notes:** Authors contributed equally (Elisa Gobbini, Margaux Hubert). Co-senior authors (Julie Caramel, Jenny Valladeau-Guilemond).

## Abstract

**Background:** Dendritic cells (DCs) are promising targets for cancer immunotherapies owing to their central role in the initiation and the control of immune responses. Their functions encompass a wide range of mechanisms mediated by different DC subsets. Several studies have identified human tumor-associated DC (TA-DC) populations through limited marker-based technologies, such as immunostaining or flow cytometry. However, tumor infiltration, spatial organization and specific functions in response to immunotherapy of each DC subset remain to be defined.

**Methods:** Here, we implemented a multiplexed immunofluorescence analysis pipeline coupled with bio-informatic analyses to decipher the tumor DC landscape and its spatial organization within melanoma patients’ lesions, and its association with patients’ response to immune checkpoint inhibitors (ICI). For this aim, we analyze a cohort of 41 advanced melanoma patients treated with anti- PD1 alone or associated with anti-CTLA4. Distance and cell network analyses were performed to gain further insight into the spatial organization of tumor-associated DCs. A Digital Spatial Profiling analysis further characterized ecosystem of tumor-infiltrating DCs.

**Results:** Plasmacytoid DCs (pDCs) were the most abundant DC population, followed by conventional cDC1 and mature DCs, present in equal proportions. In contrast to CD8^+^ T cell frequency, and despite varying densities, all DC subsets were associated with a favorable response to ICI. Distance and cell network analyses demonstrated that tumor-infiltrating DCs were largely organized in dense areas with high homotypic connections, except for cDC1 that exhibited a more scattered distribution. We identified four patterns of ecosystems with distinct preferential interactions between DC subsets. Significantly, the proximity and interactions between CD8^+^ T cells and cDC1 were positively associated with patients’ response to ICI.

**Conclusions:** Our study unravels the complex spatial organization of DC subsets and their interactions in melanoma patient lesions, shedding light on their pivotal role in shaping the response to ICI. Our discoveries regarding the spatial arrangement of cDC1, especially with CD8+ T cells, provide valuable clues for improving immunotherapeutic strategies in melanoma patients.

**What is already known on this topic:** Dendritic cells (DCs) are promising targets for cancer immunotherapies owing to their central role in the initiation and the control of immune responses. Although conventional type 1 dendritic cells (cDC1) were proposed to contribute to immunotherapy response, their precise functions and interactions with other immune populations in human cancers are largely unknown.

**What this study adds:** This study provides a precise characterization of the spatial distribution and organization of tumor-infiltrating DCs in a large cohort of advanced melanoma patients, and in correlation with response to immunotherapy. While DCs are organized in dense areas with high homotypic connections, cDC1 exhibit a more scattered distribution and form heterotypic aggregates with other DC subsets. More importantly, a close connection between cDC1 and CD8 T cell is uniquely correlated with the patients’ response to immunotherapy.

**How this study might affect research, practice or policy:** This study improves our understanding of CD8-DC spatial organization within the tumor microenvironment and will have a broad spectrum of implications in the design of anti-tumor immune-activating compounds and the design of biomarkers of response to immunotherapy for melanoma patients.

## Introduction

The incidence of cutaneous melanoma has steadily increased over the last decades and its mortality rate is the highest for skin cancers, with a 5-year survival rate of 29.8% in metastatic patients. Immunotherapy has radically shifted the treatment strategy for melanoma patients, with anti-PD-1 monotherapy or its combination with anti-CTLA4 as first-line treatment for stage IV metastatic patients, regardless of BRAF mutational status (1). However, the number of patients displaying durable response remains low due to primary or acquired resistance to immune checkpoint inhibitors (ICI) (2). The tumor immune microenvironment (TIME) plays a key role in tumor immune surveillance and immune evasion, thus contributing to the efficacy of immunotherapy (3). Although dendritic cells (DCs) represent a very small proportion of leucocytes, they play a central role in the immune response against cancer. As antigen (Ag) presenting cells (APCs), they have a unique capacity to prime naïve T cells, being able to link innate and adaptive immunity (4,5). The distinction between plasmacytoid (pDCs) and myeloid/conventional (cDCs) lineages is widely accepted and is conserved in blood and tissues (6,7). pDCs are characterized with the specific surface markers CD123 and BDCA2. In cancer, pDCs may have a dual role, as they can activate the anti-tumor response in an innate or adaptive manner (8,9). Still, their IFN-I production can be impaired by soluble factors such as TGF-β and TNF-α produced by the tumor microenvironment, thus driving regulatory T cells (Tregs) expansion (10). Type 1 conventional dendritic cells (cDC1) are characterized by the unique expression of CLEC9A and XCR1, respectively involved in endocytosis of necrotic material and in the cross-talk with T and natural killer (NK) cells (11–13). cDC1 are also particularly effective in Ag cross-presentation (11,14,15). cDC2 is a more heterogenous population able to induce Th1 but also Th2 and Th17 immune responses (16). Initially classified as DC due to their migratory and APC capacity, Langerhans cells (LCs) rather belong to the macrophage family due to their ontogeny although they share most surface markers with cDC2. Their localization is restricted to stratified epithelia, they express high levels of CD207, CD1a and EpCAM and have dichotomic roles from tolerance to Th2 polarization (17). Finally, mature DCs expressing high levels of activation markers such as DC-LAMP and CCR7, are well-conserved across cancer tissues (18), and can engage in crosstalk with T lymphocytes and NK cells through IL-12 and IFN- γ (19,20).

Some studies have investigated the prognostic impact of DC subsets across different tumor types including melanoma. While conflicting results were reported for pDCs and cDC2 (depending on the detection method e.g. flow cytometry *versus* signatures from RNA-seq)(21), the association between cDC1 and a favorable prognosis is not debated, as evidenced by our team and others (22–25). A strong infiltration of mature DCs was reported in association with improved survival and reduced metastatic spread in melanoma patients (26). However, little is known about the relative frequency? and spatial distribution of DC subsets *in situ* in tumors and their predictive value for patient survival and/or response to immunotherapy. Indeed, few spatially-resolved studies have reported DC subset infiltration in patients, and most of these analyzed a limited portion of tumor sample (e.g. often TMA), thus overlooking important information about their distribution and heterogeneity across tumor compartments, likely providing a biased analysis of rare DC subsets.

Here, we dissected the spatial organization of DC subsets of cutaneous lesions from advanced melanoma patients according to their clinical response to immunotherapy using a multiplexed immunofluorescence analysis of the whole tumor tissue. We demonstrated that pDCs are the most abundant DC population, followed by conventional cDC1 and mature DCs, which are equally represented. Despite varying densities, all DC subsets were associated with a favorable response to ICI. Interestingly, tumor-associated DCs (TA-DC) formed various clusters giving rise to four distinct spatial ecosystems composed of pDC only, mature DC only or mixed aggregates including pDCs/cDC1 or pDCs/mature DCs. Finally, we unveiled that cellular interactions between cDC1 and CD8+ T cells were strongly associated with a positive response to immunotherapy in melanoma patients.

## Results

### Several DC subsets simultaneously infiltrate human cutaneous melanoma lesions

In order to clarify the role of DC populations in the response to immune checkpoint inhibitors (ICI) in human, we analyzed their *in situ* distribution in 41 advanced melanoma patients (MELPREDICT cohort) from whom we had collected formalin-fixed paraffin-embedded (FFPE) cutaneous lesions at diagnosis. Clinical characteristics of the cohort are summarized in **Fig. 1**. All patients received a first-line treatment with ICI (anti-PD-1 monotherapy or in combination with anti-CTLA-4) as per standard of care at the Lyon Sud Hospital (Lyon, France). Patient skin biopsies of primary sites (n=20, 49%) or metastases (n=21, 51%) were performed prior immunotherapy onset. An automated seven-color multiplex immunofluorescence staining was set up to identify DC subsets **(Fig. 2A)**, using DAPI staining to perform cell annotation, BDCA2 (also named CLEC4C/CD303) for pDC detection, DC-LAMP (LAMP3) for mature DCs and CD1a for LCs. The cDC1 subset was detected using XCR1, the specificity of which was previously validated using coupled staining of CLEC9A mRNA by *in situ* hybridization and XCR1 by immunofluorescence (27). As cDC2 represent a very heterogenous population with many markers shared with macrophages, we couldn’t include this subset in the present analysis as we would need at least ten markers to identify them properly. CD8 and SOX10 were used to stain cytotoxic T lymphocytes and melanoma cells, respectively. Tissues were annotated by a pathologist to select the whole tumor area. Tissue and cell segmentation, marker quantification, as well as cell phenotyping were performed using Inform software **(Supp. Fig. 1A)**. We confirmed that each marker was specifically expressed by the corresponding cell population, thus validating the algorithm used for cell phenotyping **(Supp. Fig. 1B)**. Whole tissue sections, representing a median total surface of analyzed tissue of 27.5 mm^2^ (+/− 78.9 mm^2^, about 50-fold larger than a TMA, **Supp. Fig. 1C**), were then analyzed to decipher the spatial organization. We conducted a bioinformatic 2D reconstitution of the tissue using the spatial coordinates of the annotated cells, and obtained a readout of the cellular organization of 31 whole tumors **(Supp. Fig. 1D)**. Of note, LCs were never observed in tumor nests or tumor-associated stroma, but were only located in adjacent healthy skin, and were therefore excluded from subsequent analyses **(Fig. 2A)**.

**Figure 1:**
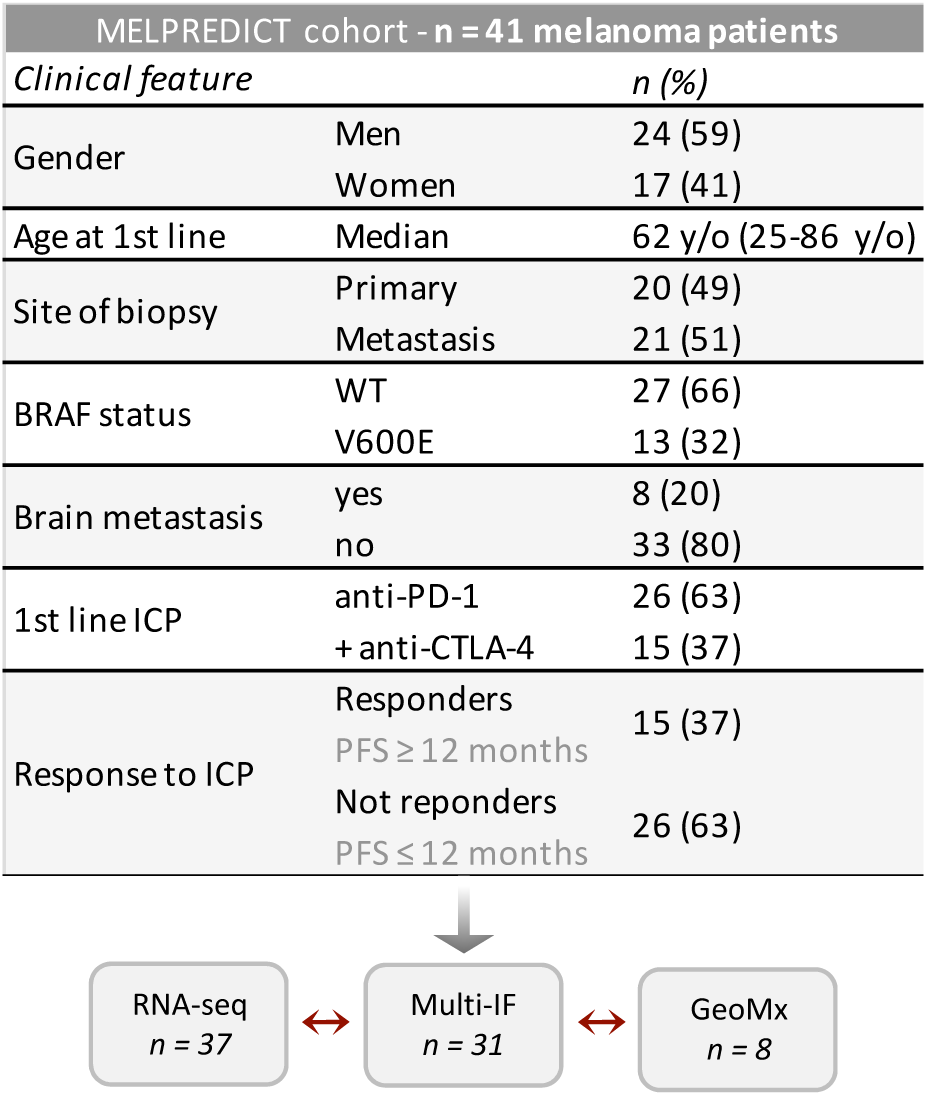
Study design and description of the MELPREDICT cohort.

**Figure 2:**
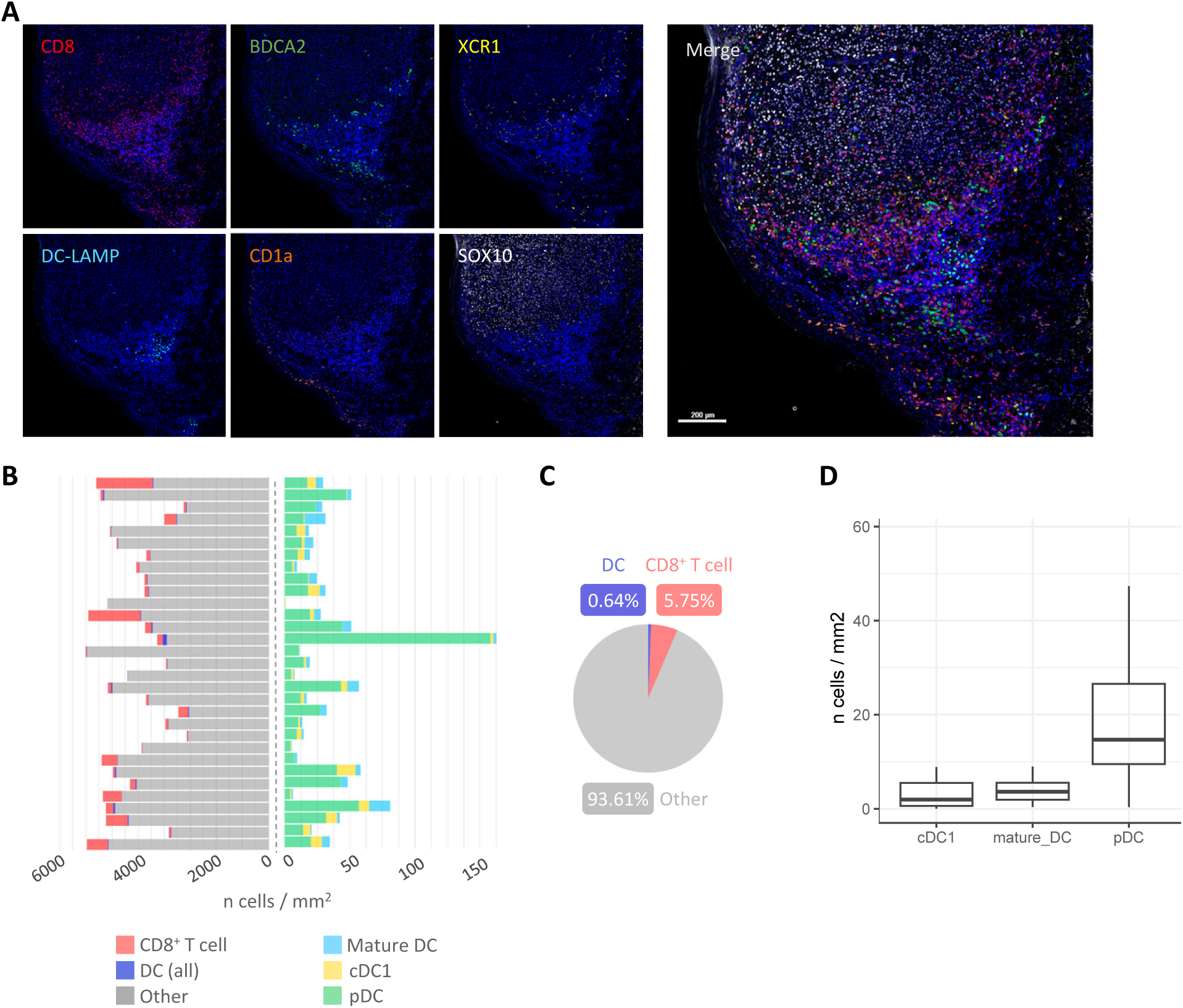
Four DC subsets can be visualized simultaneously in human melanoma by *in situ* multi- immunofluorescence. (A) Representative example of a melanoma sample stained by a multi-IF panel including CD8 (CD8 T cells), BDCA2 (pDCs), XCR1 (cDC1), DC-LAMP (mature DCs), CD1a (LCs) and SOX10 (tumor cells). (B) Cell densities in all patients. Each row represents a sample. Left panel represents densities of total DC, CD8 T cells and cells annotated as “Other” (all other cells). Right panel represents DC subset densities. (C) Median of cell proportions in the whole cohort for cells annotated as DC (all subsets), CD8 T cells and “Other”. (D) Boxplot of cell densities of all DC subsets in the whole cohort.

A quantitative analysis of whole tumors revealed that total DCs represented 0.64% (+/− 0.67%) of all annotated cells (median of 145,886 total cells) **(Fig 2B-C)**. We detected a median value of 24.7 DCs/mm^2^, ranging from 0.77 DCs/mm^2^ to 162.6 DCs/mm^2^ across samples, highlighting a consequent inter-patient heterogeneity **(Fig 2B)**. The pDC subset was the most abundant, accounting for 70.93% of all DCs, with a median density of 14.73 cell/mm^2^ (+/− 28.9 cells/mm^2^) **(Fig. 2D and Supp. Fig 1E)**. Interestingly, we observed that cDC1 were as abundant as mature DCs (14.88% and 14.80% of all DCs, respectively) with a median of 4.24 mature DCs/mm^2^ (+/− 3.9 cells/mm^2^) and 3.28 cDC1/mm^2^ (+/− 3.5 cells/mm^2^) **(Fig. 2D)**. Hence, our results highlight for the first time the *in situ* localization of DC subsets in whole samples of cutaneous melanoma and reveal that pDC are the most abundant whereas cDC1 are just as many as matDC.

### While cDC1 are spatially dispersed, mature DCs and pDCs form clusters in melanoma lesions

To decipher the spatial organization of each DC subset, we first analyzed their homotypic aggregation using the k-nearest neighbors’ algorithm (kNN), and observed that the homotypic distance between cDC1 cells was significantly greater than that between pDCs and between mature DCs **(Fig. 3A)**. Hence, cDC1 displayed a scattered distribution compared to mature DCs and pDCs; the latter being the most aggregated subset. However, we visually noticed that mature DCs were less dispersed than other DC subsets and more organized in patches. To rule out the bias due to the higher density of pDCs compared to other TA-DC populations, we used the SPIAT R package to define homotypic aggregates of DC subsets. Briefly, SPIAT defined DC cell clusters as cells from the same subset within a circle of 100μm from each other and composed of at least 10 pDCs or mature DCs **(Fig. 3B)**. To estimate the area of each aggregate, we calculated the perimeter of a convex hull defined by cells in the outermost layer of each DC aggregate. We observed that mature DC aggregates are smaller in size and further apart than pDC aggregates that remain closer together **(Fig 3C-D).** These results thus illustrate a preferential spatial organization of the distinct DC subsets *in situ*, suggesting their different modes of action to regulate the immune response.

**Figure 3:**
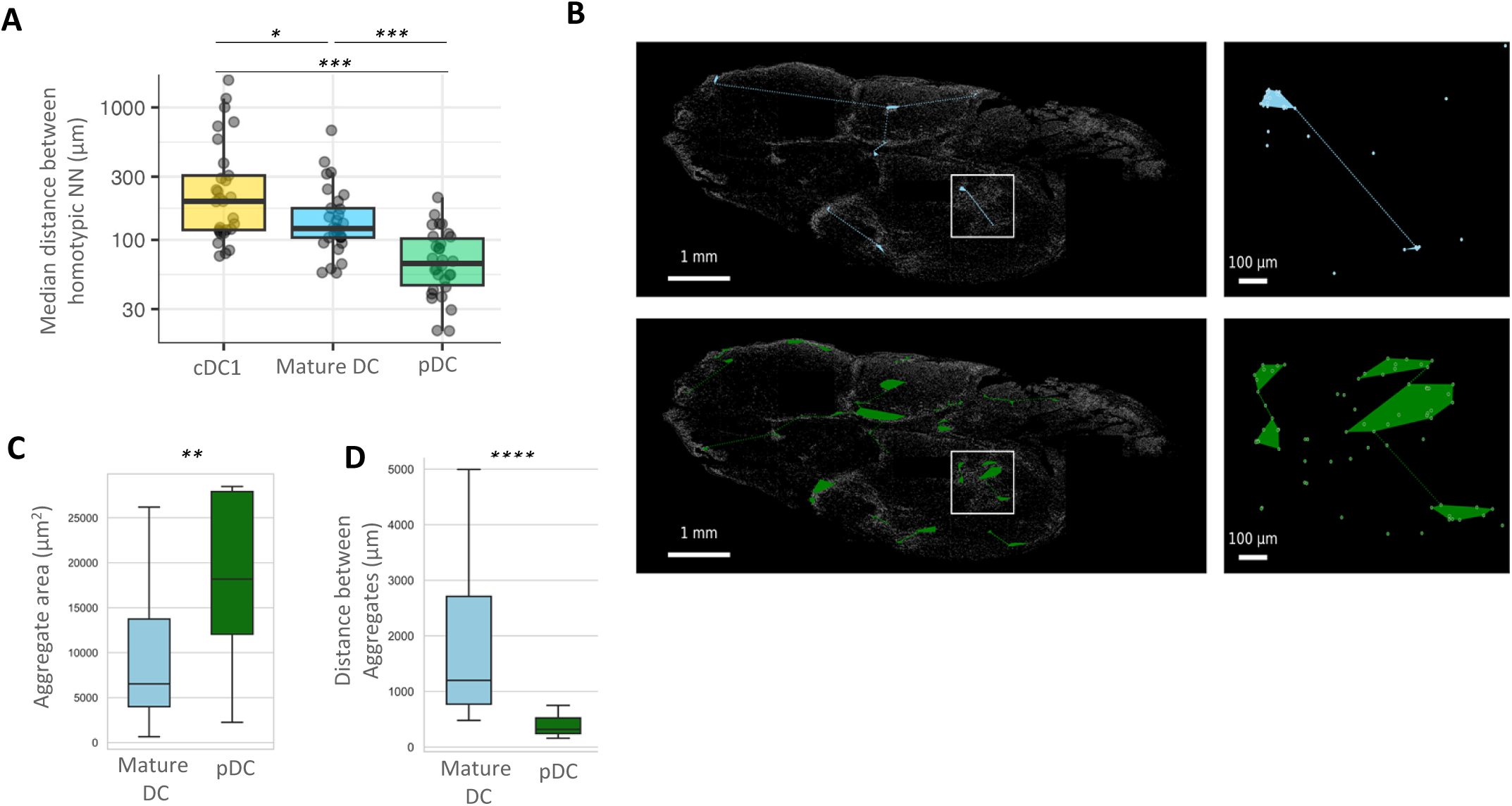
While cDC1 are mostly isolated, mature DC and pDC organize into homotypic aggregates. (A) Nearest Neighbors analysis giving the median minimal distance between DCs belonging to the same subset. (B) Example of pDC (green) and mature DC (light blue) homotypic aggregates identified with the SPIAT R package in a melanoma sample. Colored lines described the distance between aggregates of the same type calculated with the Scypy Python package. (C) Median areas of mature DC and pDC aggregates in the whole cohort calculated with the Scypy Python package. (D) Median distance between matures DC and pDC aggregates in the whole cohort calculated with the Scypy Python package. Group comparison performed by the Mann-Whitney test. * p value ≤ 0.05; ** p value ≤ 0.01; ***p value ≤ 0.001.

### Melanoma-infiltrating DC subpopulations spatially organize into four different ecosystems

As interactions and cell cooperation between DC subsets could be crucial in cancer immunity (28), we thus analyzed the co-localization and spatial organization of all DC subsets to identify heterotypic DC aggregates using the SPIAT tool. Interestingly, 30% of cDC1 were isolated cells and less than 45% belonged to DC aggregates (>10 DCs) **(Fig. 4A)**. In contrast, pDCs were mostly present in high density aggregates **(Fig. 4A)**. Next, we selected the 805 heterotypic aggregates identified across the 31 patients and performed an unsupervised hierarchical clustering based on the frequency of each subset within each aggregate to identify preferential co-localizations between DC subsets defining distinct ecosystems **(Fig 4B**, **Supp. Fig. 2**). Using this strategy, we unveiled 6 distinct clusters of DC aggregates: a large cluster mainly enriched in pDCs (C6), a cluster mainly enriched in mature DCs (C2), 2 clusters with a mixed composition of cDC1 and pDC (C3-C5), 1 cluster with a mixed composition of mature DCs and pDCs (C1), and one cluster composed of only 2 aggregates belonging to the same tumor and with a mixed composition between cDC1 and mature DCs (C4). Interestingly, we did not identify aggregates enriched specifically in cDC1, consistent with their scattered distribution evidenced above. This analysis unveiled that DC subsets were spatially organized into four main ecosystems; two of these mainly displayed a homotypic composition, enriched in either pDCs or mature DCs, and two heterotypic ecosystems composed of pDCs and cDC1 (purple), or of pDCs and mature DCs (green) **(Fig 4D down)**.

**Figure 4.**
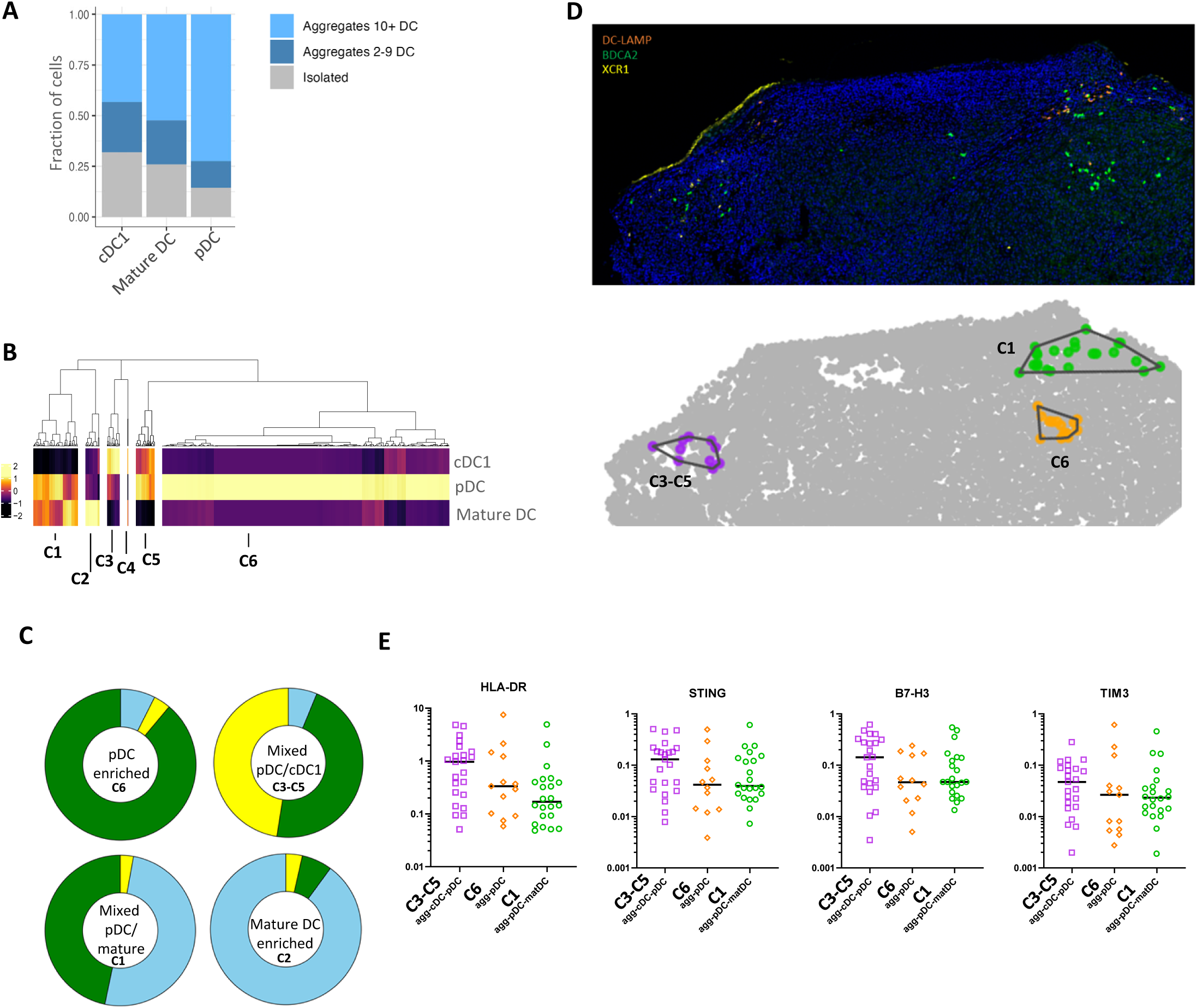
Four patterns of heterotypic DC ecosystems can be defined in melanoma. (A) Proportion of aggregated / isolated cell for each DC subset. (B) Unsupervised hierarchical clustering of DC aggregates according to their DC-composition (% of pDC, cDC1 and mature DC). Scale represents the Z score. (C) Proportion of each DC subsets within the 4 patterns of DC ecosystems. (D) Example of 3 defined DC ecosystems generated with SPIAT and visualized on multi-IF image (top panel) or annotated on 2D reconstitution (bottom panel). (E) DSP analysis made on homotypic pDC aggregates (n=13 ROIs), and heterotypic pDC aggregates (with matDC n=22 ROIs and with cDC1 n = 23 ROIs).

To gain insight into variations in the tumor microenvironment (TME) surrounding the distinct DC ecosystems, we performed high-plexed spatial protein analysis (GeoMx® Digital Space Profiler, Nanostring^©^) on 8 patients, using a panel consisting of 32 markers, encompassing the various immune populations and immune checkpoints.

Given that DC-LAMP is frequently used to characterize TLS, we assumed that mature DC-enriched aggregates (detected using the same marker) corresponded to tertiary lymphoid structures (TLS) (29). We thus focused on the three other ecosystems that are undescribed to date. Interestingly, pDC-cDC1 aggregates expressed high levels of HLA-DR, STING and B7-H3 levels, suggesting a crosstalk between these two subsets leading to their mutual activation. Moreover, this ecosystem also expressed the highest level of TIM3 **(Fig 4E).** This receptor is present on memory or activated T lymphocytes (LT) (Th1 LT-CD4, regulatory LT [Treg] and cytotoxic LT-CD8^+^ [Tc1]), B lymphocytes, NK cells, but also myeloid cells including DCs and thus suggesting an immune activation associated with pDC-cDC1 aggregates **(Fig 4E).**

### The density of DC subpopulations and their spatial organization are crucial for melanoma patient’s response to ICI

We then assessed the association between each DC subsets’ density/spatial organization and the response to immunotherapy. Patients were stratified as responding (R, n=12) or non-responding (NR, n=19) according to their progression-free-survival (PFS) evaluated at 1-year following onset of first-line immunotherapy **(Fig. 1)**. As shown in Supplementary Table 1, there is no confounding parameter associated with the response. Interestingly, the density of each subset, namely cDC1, mature DCs and pDCs was significantly higher in responding patients (**Fig. 5A**). Total DC aggregates were also enriched in responding patients **(Fig. 5B)**, supporting our results on the preferential spatial organization pattern of mature DCs and pDCs as aggregates. Interestingly, responding patients displayed DC subsets closer together both in the case of pDCs and mature DCs, suggesting that homotypic organization and closer interactions between pDCs or mature DCs are essential for the efficacy of immunotherapy (**Fig. 5C-E**). Conversely, the median distance separating cDC1 was not associated with response to ICI, suggesting that cDC1 may spatially interact with other immune cells to regulate the tumor immune response (**Fig. 5E-F**).

**Figure 5:**
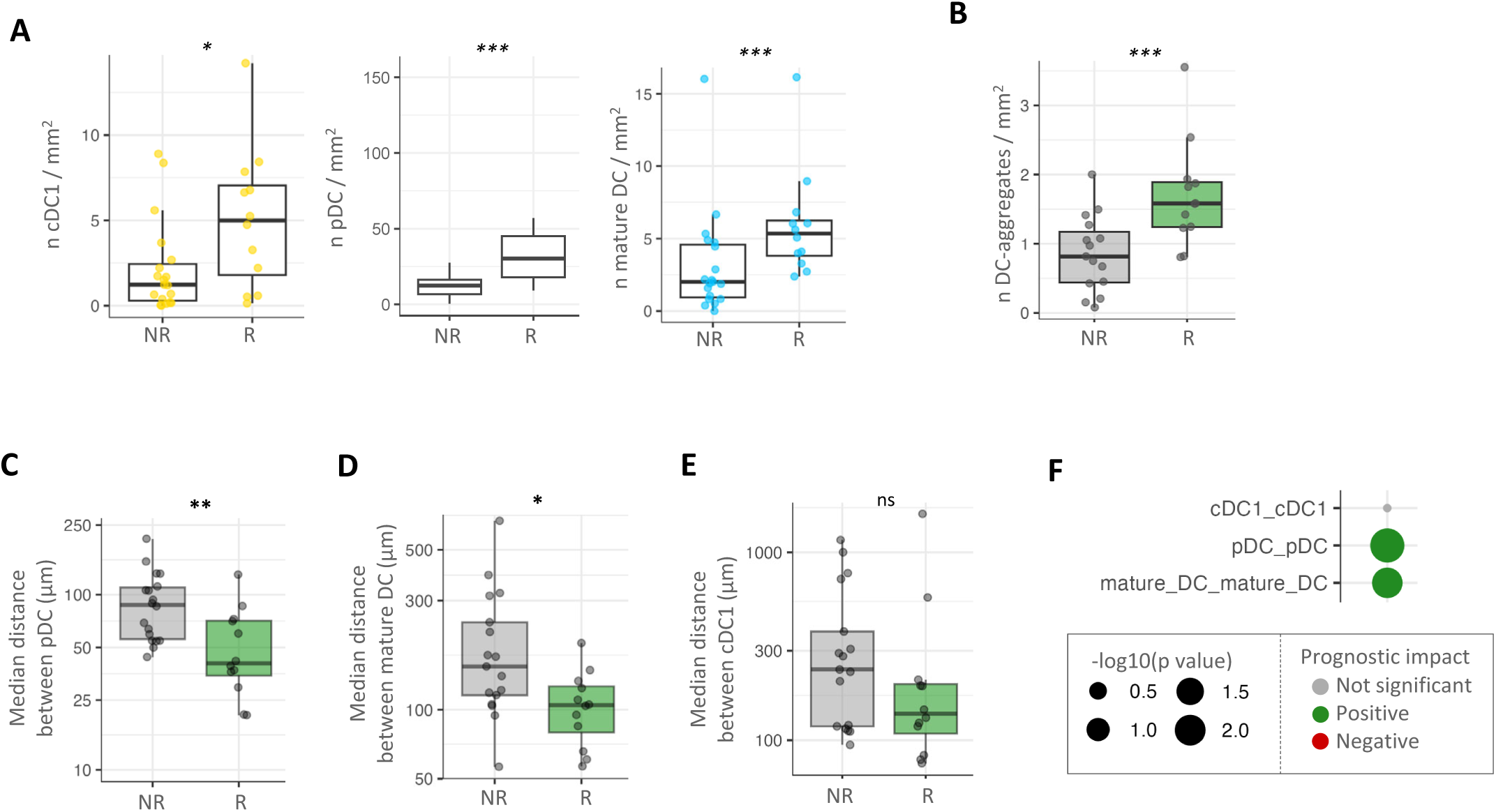
DC subsets and their spatial organization are associated to the ICI response. (A) Comparison of cDC1, pDC and mature DC densities between responder (n=12 patients) and not responders (n=19 patients). (B) Comparison of DC aggregate number in responder and non-responder patients. (C) Nearest Neighbors analysis measuring the median minimal distance between pDC in responder and not responders. (D) Nearest Neighbors analysis measuring the median minimal distance between mature DC in responder and not responders. (E) Nearest Neighbors analysis measuring the median minimal distance between cDC1 in responder and not responders. (F) Bubble plot illustrating the p-value referring to the comparison between responders and not responders for the median minimal distance among cells annotated in the same DC subset. Group comparison performed by the Mann-Whitney test. * p value ≤ 0.05; ** p value ≤ 0.01; ***p value ≤ 0.001.

### Spatial organization between cDC1 and CD8^+^ T cells is pivotal for response to ICI in melanoma patients

As DCs are instrumental in activating cytotoxic T cells, we next evaluated the spatial interconnections between DC subsets and intra-tumoral CD8^+^ T cells according patients’ response to immunotherapy. We observed that the CD8^+^ T cell density did not differ between responding and non-responding patients (**Fig. 6A**). We confirmed these results by scoring the CD8^+^ T cell signature using MCP counter from our RNA-seq data of the cohort that we previously published (30) **(Fig. 6B)**. We then focused on CD8^+^ T cell distribution by using the kNN algorithm and observed that a shorter distance between CD8^+^ T cells was associated with a better response to immunotherapy (**Fig. 6C**). To gain further insight into the spatial architecture of CD8^+^ T cells, we performed a Delaunay triangulation to reconstitute networks between all cells **(Fig. 6D)**. Interestingly, the number of links between CD8^+^ T cells was significantly higher in responders **(Fig. 6E)**. Together, these results suggest that the structure of the CD8^+^ T cell network, rather than its density, is required for the efficacy of immunotherapy. Given that the DC composition in the TME influences CD8^+^ T cell infiltration and activation (31,32), we then analyzed the correlation and spatial organization between CD8^+^ T cells and DC subsets. The correlation between CD8^+^ T cell density and mature DCs was greater than that with other DC subsets **(Fig. 6F)**. Interestingly, although we found a similar distance between all DC subsets and their closest CD8^+^ T cell neighbor **(Fig. 6G)**, the median distance between CD8^+^ T cells and cDC1 was markedly reduced in responding patients (**Fig. 6H**). Consistently, we observed a greater number of connections between CD8^+^ T cells and cDC1 in responding patients (**Fig. 6I**). In contrast, the number of links between CD8^+^ T cells and mature DCs or pDCs was not associated with response to ICI **(Fig. 6J)**. Based on these results, we believe that the spatial organization of CD8^+^ T cells, in particular their crosstalk with cDC1, is important and could be predictive of patient response to ICI in melanoma.

**Figure 6:**
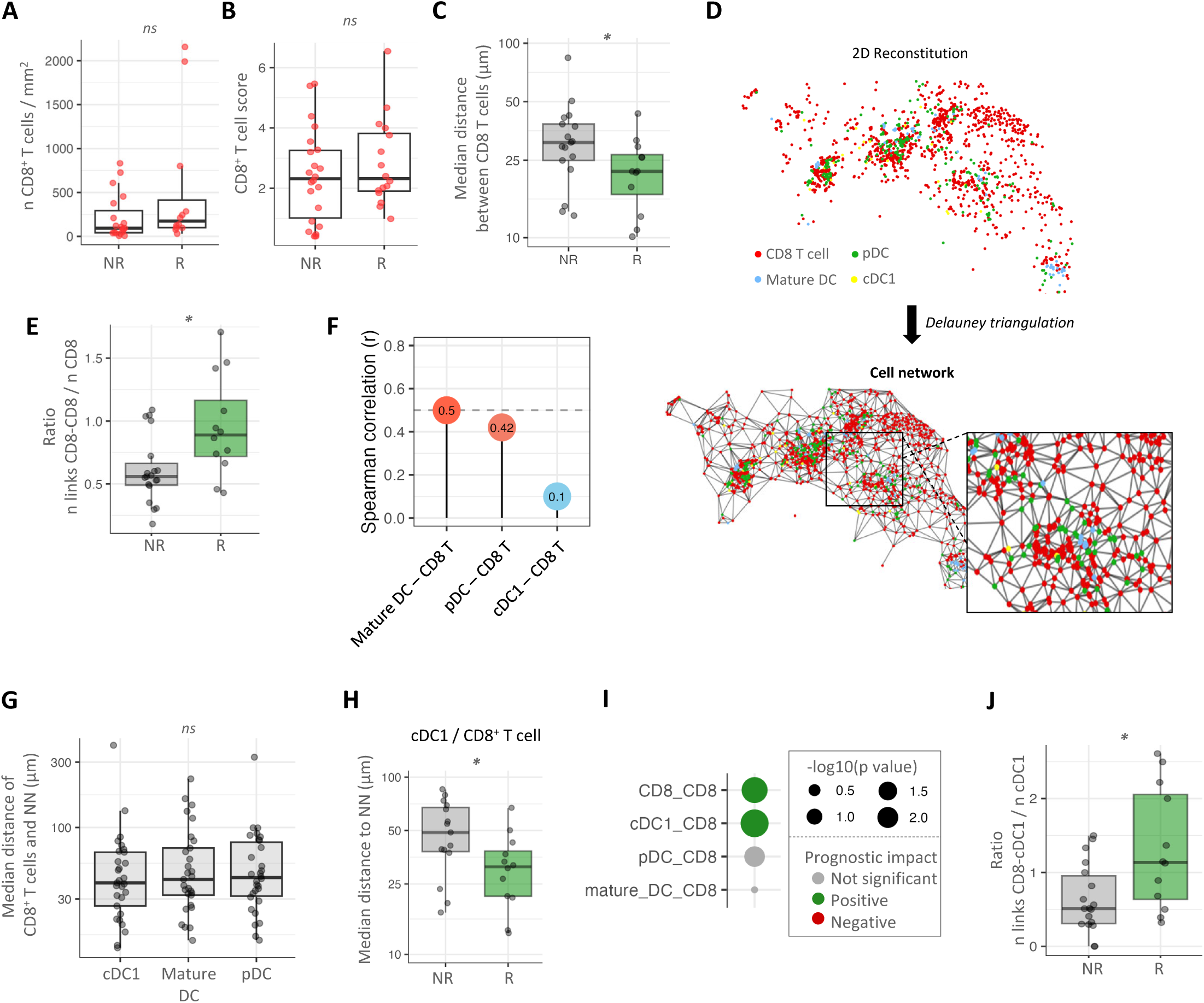
cDC1 spatial organization with CD8 T cell is associated with the response to ICI. (A) Comparison of CD8 T cells density between responders and not responders. (B) CD8 T cell signature score in responder and not responders obtained with MCP-counter using RNA- seq expression data. (C) Comparison of the median distance separating CD8 T cells in responders and non responders using a kNN analysis. (D) Example of the cell network created by the Delaunay triangulation from the spatial coordinates of annotated cells on a melanoma sample. Outlier links > 200 μm were filtered out. (E) Ratio between the number of connections between DC /CD8 T cells and the total number of DC calculated by the Delaunay triangulation on the whole cohort in responder and non-responders. (F) Spearman correlation between annotated cells densities in the whole cohort. “r” correlation coefficient are reported in the bubbles. (G) Median minimal distance separating CD8 T cells from cDC1, pDC and mature DC, generated by nearest neighbors analysis performed with DC as reference cells. (H) Comparison of the median minimal distance separating CD8 T cells from cDC1 between responders and non responders, generated by nearest neighbors analysis performed with cDC1 as reference cells. (I) Bubble plot illustrating the p value of the median distance between nearest neighbors considering DC subsets as reference and CD8 T cells as neighbors, compared between responders and not responders. Group comparison performed by the Mann-Whitney test. * p value ≤ 0.05; ** p value ≤ 0.01; ***p value ≤ 0.001. (J) Ratio between the number of connection cDC1- CD8 T cells and the total number of cDC1 calculated by the Delaunay triangulation on the whole cohort in responder and not responders.

## Discussion

Dendritic cells play a critical role in orchestrating and shaping the immune response. Despite previous studies demonstrating the infiltration of pDCs, cDC1, cDC2, LCs and mature DCs in dilacerated melanoma samples (22), the *in situ* spatial organization of DCs and their impact on prognosis and response to immunotherapy remains largely unknown. Moreover, previous studies describing tumor-associated DC subsets by *in situ* approaches focused on up to two subsets at once and worked on small areas of tumor tissue (26,33–35). It precluded capturing the complexity and heterogeneity of their spatial distribution in tumors. In this study we demonstrated for the first time the simultaneous presence of pDCs, cDC1, LCs and mature DCs in cutaneous melanoma patient lesions *by in situ* multiplexed immunofluorescence and the importance of the cDC1-CD8 interaction to favor the response to immunotherapy.

LCs in skin cancers were reported to contribute to skin inflammation and to tumor immune evasion in melanoma (36). We observed that LCs were always excluded from tumor nests or tumor-associated stroma but were exclusively present in healthy skin. Neagu *et al.* showed that CD1a+ LC were either absent or scarce within the tumor mass and were associated with good prognostic features such as thinner and not ulcerated tumors (37). This suggests a limited involvement of LCs in the anti-tumor response and corroborates our recent findings on TCGA public data, querying the prognostic impact of LCs using a LC gene signature, and showing no impact on survival, including for melanoma patients (23). This also agrees with Howell *et al.,* who demonstrated that melanomas fail to activate LC migration to lymph nodes until tumors reach a critical size (38).

Here, for the first time, cDC1 were detected *in situ* to analyze their localization and their spatial organization compared to other DC subsets. In mice, cDC1 are essential for the induction of anti-tumor immunity and responses to immunotherapies. In a conditional PD-L1 knock-out model on cDC1, Oh *et al.* reported that, despite PD-L1 being expressed on most myeloid cells, inhibiting PD-L1 on cDC1 was sufficient to inhibit tumor growth by increasing the infiltration of CD8^+^ T cells with an activated phenotype (PD1^+^, LAG-3^+^, TIM-3^+^) (39). Moreover, while the treatment of WT mice with a combination of immune checkpoints blockade (anti-CD137 + anti-PD-L1) led to the rejection of B16 melanomas, Batf3^-/-^ mice lacking cDC1 did not respond to this combined immunotherapy (40). In the same tumor model, cDC1 were also essential for the response to anti-PD-L1 combined with a BRAF inhibitor (41,42). In human samples, we and others have shown that a high cDC1 infiltration score is associated with favorable patient prognosis (13,23,25). Moreover, Barry *et al*. demonstrated the association between cDC1 signature enrichment and improved survival in patients with metastatic melanoma. They showed that cDC1 were more abundant in melanoma patients responding to first-line immunotherapy compared to non-responding patients by flow cytometry on dissociated tumors (43). However, cytometry and transcriptomic analysis obviously do not provide information about DC subset localization *in situ*, their interaction with other immune populations and how this could impact the response to immunotherapy.

The spatial organization of CD8^+^ T cells rather than their abundance determines the response to immune checkpoint inhibitors. Indeed, we observed here that in responding patients, CD8^+^ T cells formed dense compartments with strong homotypic interactions. In contrast, CD8+ T cells presented a more diffuse distribution in non-responding patients. Similar results were found by Xiao *et al.* using imaging mass cytometry on melanoma tissue (44). However, that analysis did not focus on CD8^+^ T cells but included all lymphocytes (B cells, CD4^+^ T cells, CD8^+^ T cells, double-positive T cells and Treg) and did not provide information about the co-localization with DC subsets. The different spatial relationships between DCs and CD8^+^ T cells suggest the existence of specificities in driving the anti-tumor immune response according to the DC subset involved.

The mechanisms underlying the impact of cDC1 on patient outcome are largely unknown. The ability of cDC1 to produce large amounts of CXCL9/10 and their expression of XCR1 are two properties that suggest a close cross-talk with immune effector cells involved in a cytotoxic response. It has been previously shown that cDC1 have a greater capacity to activate cytotoxic immune responses through antigen cross-presentation compared to other DC subsets (14,45–47). Moreover, the bidirectional cross-talk between DCs and NK cells has been elucidated in several studies (15,48). While DCs can activate NK cells, these latter can in turn affect the recruitment and maturation of DCs via FLT3 and IFN-γ production. In this work, we unveiled no correlation between CD8^+^ T cell and cDC1 infiltration. However, in patients responding to immunotherapy, cDC1 were closer to CD8^+^ T cells than in non-responding patients. Of note, no difference in proximity between pDCs or mature DCs and CD8^+^ T cells was found according to the response to immunotherapy. This supports the idea of a privileged interaction between cytotoxic lymphocytes and cDC1 to orchestrate the anti-tumor response in immunotherapy-treated patients as it was recently shown in melanoma mouse models by Meiser *et al.* and Magen *et al.* (32,49). Antoranz *et al.* recently demonstrated that the phenotype and function of CD8^+^ T cells varied according to their localization, closer to the stroma-tumor interface or to the center of the melanoma tissue (50). In addition, CD8^+^ T cells outside the tumor are enriched in TCF1/TCF7 transcription factors (51) with a weak expression of co-inhibitor molecules, representing a functional population with likely anti-tumor activity. Conversely, CD8^+^ T cells closer to the tumor edge and infiltrating the tumor, are more likely characterized by a progressively higher expression of co-stimulatory molecules such as PD-1, TIM-3 and VISTA, acquiring a more exhausted and dysfunctional phenotype (52). Investigating the phenotypes and functional characteristics of CD8^+^ T cells interacting with cDC1 would be informative on their contribution to activate an anti-tumor cytotoxic response.

Moreover, based on our spatial analysis of DC aggregates, we demonstrated that patients with tumors enriched in pDC aggregates were more likely to respond to immunotherapy. Previous studies alternatively reported a negative or a positive prognostic impact of pDCs in cancer (22). pDCs can directly attack tumor cells in a TNF-related apoptosis-inducing ligand (TRAIL)-dependent manner or can contribute to an immunosuppressive microenvironment, priming cells involved in IL-10 production such as Tregs, via the ICOS-ICOSL axis stimulation (53). Some receptors uniquely expressed by human pDCs, such as ILT7 and NKp44 can also dampen IFN-α production, leading to Treg expansion and could favor tumor immune escape (10). Recent work also suggested that pDCs might induce and regulate tumor-directed T cell responses within TLS (54). pDCs have been described in colon cancer-associated TLS and, in line with our results, were preferentially located close to CD4 T cells within the T cell zone and frequently presented an activated phenotype (IRF7+). Here, our work unveiled aggregates in which pDCs clustered with cDC1 (pDC/cDC1 aggregates) or mature DCs (pDC/mature DC aggregates). Previous studies focusing on the localization and phenotype of TLS-associated DCs, demonstrated that a high frequency of TLS-associated DCs expressing the maturation marker LAMP3/DC-LAMP was linked to a robust infiltration of T cells (55). The density of mature DCs was also associated with expression of genes related to T-cell activation, Th1 phenotype and cytotoxic polarization. Moreover, more TLS-associated DCs was correlated with long-term survival and response to immunotherapy (56). The pDC/cDC1 enriched aggregates we described herein suggest the existence of a pDC-cDC1 crosstalk and, while this has never been confirmed in human, some evidence from infectious models supports this hypothesis. Indeed, the analysis of DC subsets within draining lymph nodes during vaccinia virus infection showed that infected cDC1 recruited pDCs via the CCR5 receptor (57). Albeit, mobilized pDCs increased the expression of maturation markers (CD80, CD86 and CD40) and antigen presentation of non-infected resident cDC1 recruited by activated CD8^+^ T cells via the XCL1/XCR1 axis. In our current study, high-plexed spatial protein analysis (GeoMx DSP) also strongly suggests this crosstalk, demonstrating higher STING, B7-H3, and HLA-DR levels in those aggregates. Finally, in cancer, a synergistic effect between pDCs and cDCs in the activation of a cytotoxic response has been suggested in a mouse model (58).

In conclusion, we implemented for the first time an *in situ* approach to characterize cDC1 infiltration in melanoma tissue along with their spatial relationship with other DC subsets and CD8^+^ T lymphocyte infiltration. Whereas all subsets were shown to be enriched in melanoma patients responding to immunotherapy, the close contact between CD8^+^ T cells and cDC1, and the organization of DCs in aggregates enriched in pDCs are particularly important characteristics linked to response to immunotherapy. Hence, cell density may not be the best criterion to evaluate to predict patients’ response to immunotherapy, unlike the spatial organization/interactions of immune cell subsets within the tumor, which seems essential for generating an efficient anti-tumor immune response.

## Materials and Methods

### Patient samples

Melanoma tumor samples were obtained from surgical specimens collected from 41 advanced melanoma patients at diagnosis in the framework of their standard of care, with written informed consent for the use of samples for research purposes obtained from all patients prior to analyses. Formalin-fixed paraffin-embedded (FFPE) cutaneous tumor samples were collected (1 slide of 4 μm) through the Biological Resource Center of the Lyon Sud Hospital (Hospices Civils de Lyon). All patients received a first-line treatment with immune checkpoint inhibitors (anti-PD1 alone or associated with anti-CTLA4) as per the standard of care. Patients were stratified as responding (R) or non-responding (NR) based on progression-free survival (PFS), measured through trimestral clinical evaluation of tumor extension by CTAP scan according to RECIST1.1 criteria, with a cut-off at one year following first-line immunotherapy initiation. No clinical parameters were found associated to the clinical response to immunotherapy in this cohort (Supp Table 1). This study was approved by a regional ethics board (Comité de Protection des Personnes Ile de France XI, Saint-Germain-en-Laye, France) for the use of clinical samples and the collection of associated clinical data according to the CNIL regulation (authorizations n°22_5680 and n°22_5896).

### Multiplex immunofluorescence staining

Markers for tumor-infiltrating DC identification were chosen based on previous literature. In particular, we selected first DAPI staining for cell segmentation, BDCA2 (Goat Polyclonal, R&D Systems, AF1376) for pDC, DC-LAMP (Rabbit 1010E1.01, Dendritics, DDX0191P) for mature DCs, XCR1 (Rabbit D2F8T, Cell Signaling, 44665S) for cDC1, CD1a (Mouse 010, DAKO, M3571) for LCs, CD8 (Mouse C8/144B, DAKO, M7103) for T cells and SOX10 (Mouse A-2, Santa Cruz, Sc-365692) for melanoma cells. We set up a sequential seven-colors multiplexed immunofluorescence by sequential staining cycles using the tyramide-signal amplification (TSA)-based OPAL seven-color Automation Research Detection Kit (Leica, DS9777) on the Bond RX automatic stainer (Leica Biosystems). Secondary antibodies used to amplify the signal included the Anti-Goat IgG HORSERADISH Peroxidase (HRP) Conjugated (Donkey Polyclonal, Life Technologies, A16005), the Anti-Rabbit IgG, HORSERADISH Peroxidase (HRP) Conjugated (Goat Polyclonal, Life Technologies, A1604) and the Anti-Rat IgG, HORSERADISH Peroxidase (HRP) Conjugated (Goat Polyclonal, Thermo Scientific, 31470). Different antigen retrieval reagent conditions, antibody concentrations, and incubation times were tested for each marker of interest on tonsil FFPE samples and melanoma. Optimized staining conditions are summarized in **Sup Table 3**.

### Dendritic cell annotations and Spatial analysis

Stained FFPE slides were scanned on a Vectra® Polaris™ Imaging System (Akoya, Biosciences, Marlborought, MA, USA) at X20 magnification. Slides were then visualized with Phenochart (1.0.12 version, AKOYA Biosciences) for multispectral imaging and were annotated and quantified using inForm Tissue Analysis Software (Akoya, Biosciences, Marlborought, MA, USA). The whole tumor area including the invasive margin, defined as the region separating host tissue from the malignant nests within a 1 mm distance, was analyzed for each sample.

For the Nearest Neighbors analysis (KNN), we used the minimum distance between each pair of cells of different or identical phenotypes using the k-Nearest Neighbors (KNN) algorithm within the sklearn.neighbors package with k=1. Using the R package SPIAT (https://github.com/TrigosTeam/SPIAT), we identified clusters of cells located within a specified distance of each other. To achieve this, we used Euclidean distances to cluster cells based on their spatial proximity. We defined cells with a 100μm radius as neighbors, and a minimum of three cells were required to constitute a cluster. Then, using the scipy.spatial module ConvexHull function (SciPy library in Python, https://docs.scipy.org/doc/scipy/reference/generated/scipy.spatial.ConvexHull.html), we computed convex hulls defined by cells in the outermost layer of aggregates. The distance between two convex hulls was calculated using the minimum distance between their closest points which was obtained by indexing the cdist (function from the module scipy.spatial.distance that computes the distance between points) matrix with the vertices attribute of the ConvexHull objects. Aggregate analysis was performed using the R package SPIAT: DC aggregates were defined as a group of DCs (min 10) included within a 100 μm radius around a reference DC. For Network and close connection analysis, we used the Delaunay Triangulation method and included all annotated cells (DC subsets phenotypes, CD8 T CELL phenotype and “other”) to generate the network using a cut-off of 30 μm to focus on close connections.

### Digital Spatial Profiling

NanoString DSP technology was used on FFPE tissue from 8 representative patients. Tissue sections were incubated with a cocktail of 32 unique oligonucleotide-conjugated antibodies and counterstained with a nuclear stain. Antibody oligos were photocleaved within the selected ROIs and collected for nCounter analysis. nCounter digital barcode counts for each antibody and ROI sample were first normalized using internal spike-in controls.

### Statistical analyses

Mann-Whitney tests for unpaired samples and Wilcoxon tests for paired samples were performed to compare two groups. To compare more groups, Kruskal-Wallis tests were performed for unpaired samples and Friedman tests for paired samples. All graphs show each sample and the median value. Statistical significance: *p<0.05, **p<0.01, ***p<0.001, ***p<0.0001.

#### Bulk-RNA-seq data and MCP counter analysis

As previously described in Plaschka *et al.* (30), the GEO accession number for this cohort is GSE169203. MCP counter was used (59) to estimate the relative abundance of several populations of immune cells.

## Acknowledgments

We would like to thank the staff of the core facilities at the Cancer Research Center of Lyon (CRCL) and the BRC (Biological Resources Center) of Lyon Sud Hospital (Hospices civils de Lyon). Elisa Gobbini was supported by ESMO (any views, opinions, findings, conclusions or recommendations expressed in this material are those solely of the authors and do not necessarily reflect those of ESMO) and ITMO INSERM. Margaux Hubert was supported by the ARC foundation. We would like also to thank our financial supports : INSERM, INCA-DGOS PRTK_2017-022 (SD, CC), INCA PLBIO INCa_4508, ANRS, ARC sign’it 2019, Ligue contre le Cancer (Régionale Auvergne-Rhône-Alpes et Saône-et-Loire, Comité de la Savoie, Comité de l’Ain), the Région Auvergne-Rhône-Alpes, SIRIC project (LYRIC, grant no. INCa_4664) and the LABEX DEVweCAN (ANR-10-LABX-0061) of the University of Lyon, within the program “Investissements d’Avenir” organized by the French National Research Agency (ANR), the Société Française de Dermatologie (JC), the association Melarnaud and Vaincre le Mélanome (JC, SD).

The authors declare that the research was conducted in the absence of any commercial or financial relationships that could be construed as a potential conflict of interest.

## Contributions

E.G designed experiments, analyzed results, did statistical analyses and wrote the manuscript. M.H. designed experiments, did bioinformatics, analyzed results, and did statistical analyses. L.H. and D.L. did bioinformatics, analyzed results, and did statistical analyses. A.C.D., C.S, S.B., P.D., L.B., J.B. and L.M. performed some experiments and analyzed results. O.H. performed the pathological annotations of samples. V.B. and M.G. provided help and advice for pathological annotations and Inform analyses. J.L. and J.P. performed DSP analysis. C.C. and B.D. provided strategic advice and revised the manuscript. S.D. and A.E. provided human samples and/or clinical data, strategic advice and revised the manuscript. J.V-G. and J.C. designed experiments, supervised the research, and wrote the manuscript.

---

**Supp Table 1:** Clinical features of MELPREDICT patients at baseline prior immunotherapy onset.

| MELPREDICT cohort |  |  |  |  |  | Test Chi-square: |
| --- | --- | --- | --- | --- | --- | --- |
| Clinical feature |  | Total (n= 41) | Analyzed (n =31) | Responders (n = 12) | Not-Responders (n =19) | P value |
| Gender | Men | 24 (59) | 18 (58) | 6 (50) | 12 (63) | .4695 |
|  | Women | 17 (41) | 13 (42) | 6 (50) | 7 (37) |  |
| Age at first line | Median | 62 y/o (25-86) | 64 y/o (35-86) | 64 y/o | 63 y/o | .4220 |
| Site of biopsy | Primary | 20 (49) | 14 (45) | 5 (42) | 9 (47) | .7560 |
|  | Metastasis | 21 (51) | 17 (55) | 7 (58) | 10 (53) |  |
| BRAF Status | WT | 27 (66) | 20 (65) | 8 (67) | 12 (63) | .5920 |
|  | V600E | 13 (32) | 10 (32) | 3 (25) | 7 (37) |  |
| Brain metastasis | Yes | 8 (20) | 6 (19) | 1 (8) | 5 (26) | .2170 |
|  | No | 33 (80) | 25 (81) | 11 (92) | 14 (74) |  |
| First line ICP | Anti-PD1 | 26 (63) | 21 (68) | 7 (58) | 14 (74) | .3731 |
|  | Anti-PD1/anti-CTLA4 | 15 (37) | 10 (32) | 5 (42) | 5 (26) |  |

**Supp Table 2:** Mann-Whitney test p-values on immune cell density depending on various clinical parameters in the MELPREDICT cohort.

| Clinical feature | CD8 | cDC1 | Mature DC | pDC |
| --- | --- | --- | --- | --- |
| Gender (M vs F) | 0.594 | 0.921 | 0.767 | 0.567 |
| Site of biopsy (Primary vs MTS) | 0.984 | 0.576 | 0.118 | 0.922 |
| BRAF status (Muté vs WT) | 0.474 | 0.948 | 0.649 | 0.100 |
| Brain MTS (Yes vs No) | 0.952 | 0.751 | 0.981 | 0.643 |
| First line ICP (Anti-PD1 vs antiPD1 + antiCTLA4) | 0.789 | 0.010 | 0.983 | 0.392 |

**Sup Table 3:** Optimized immunofluorescence staining conditions for the multiplex immunofluorescence panel.

| Marker : | Antibody Dilution | Antibody Retrieval | Antibody Incubation | Secondary Antibody |
| --- | --- | --- | --- | --- |
| DC-Lamp | 1:100 | ER2 - 20 min | 30 min | Anti-Rat IgG-HRP Conjugated |
| BDCA2 | 1:40 | ER2 - 20 min | 30 min | Anti-Goat IgG-HRP Conjugated |
| CD8 | 1:40 | ER2 - 20 min | 30 min | Opal Polymer HRP Ms + Rb |
| CD1a | 1:50 | ER2 - 20 min | 30 min | Opal Polymer HRP Ms + Rb |
| XCR1 | 1:50 | ER2 - 20 min | 30 min | Anti-Rabbit IgG-HRP Conjugated |
| SOX10 | 1:1000 | ER2 - 20 min | 30 min | Opal Polymer HRP Ms + Rb |

**Supp Fig 1:**
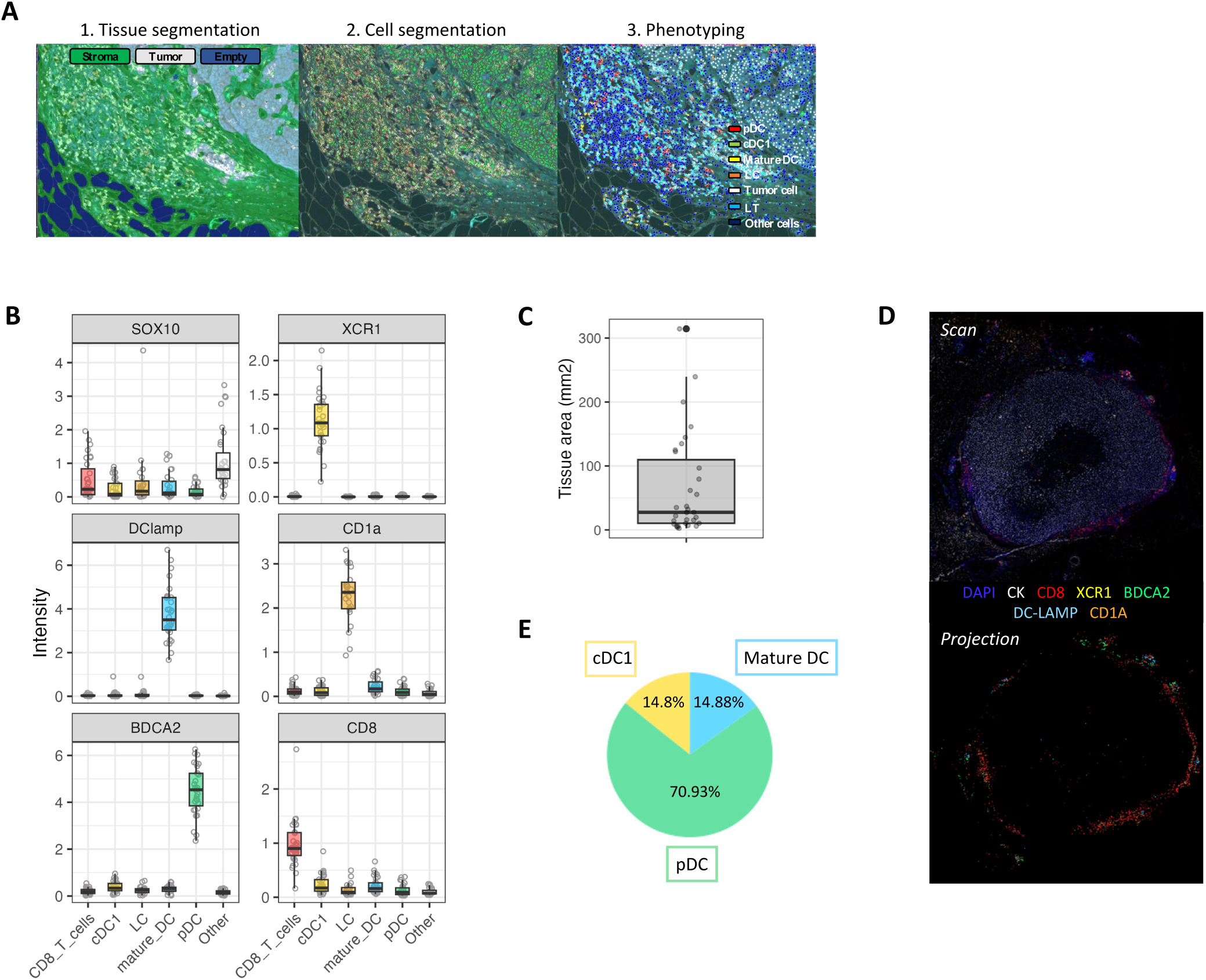
Work flow analysis for the multiplexed immunofluorescence staining. (A) Image analysis workflow performed using Inform©. Tissue segmentation (1) was trained to split up tissue (tumor and stroma) from empty/artefact zones. Cell segmentation (2) was trained to detect nuclei. Cell phenotyping (3) was trained to identify all distinct phenotypes of interest based on staining intensities and shape of cells. Results were plotted on the spectral mixed image as dots: pDCs (red), cDC1 (green), LCs (orange), mature DCs (yellow) and CD8 T cells (cyan blue). The phenotype named “Other” (blue) included all other immune and non-immune populations. (B) Specific marker expression in each annotated populations. (C) Median tissue area analyzed in the whole cohort by the multiplex immunofluorescent panel. (D) Comparison between the multi-IF image (left panel) and the 2D projection of cells using their XY coordinates and phenotype (right panel). (E) Median of DC subsets proportions among all tumor-associated DCs

**Supp. Fig. 2.**
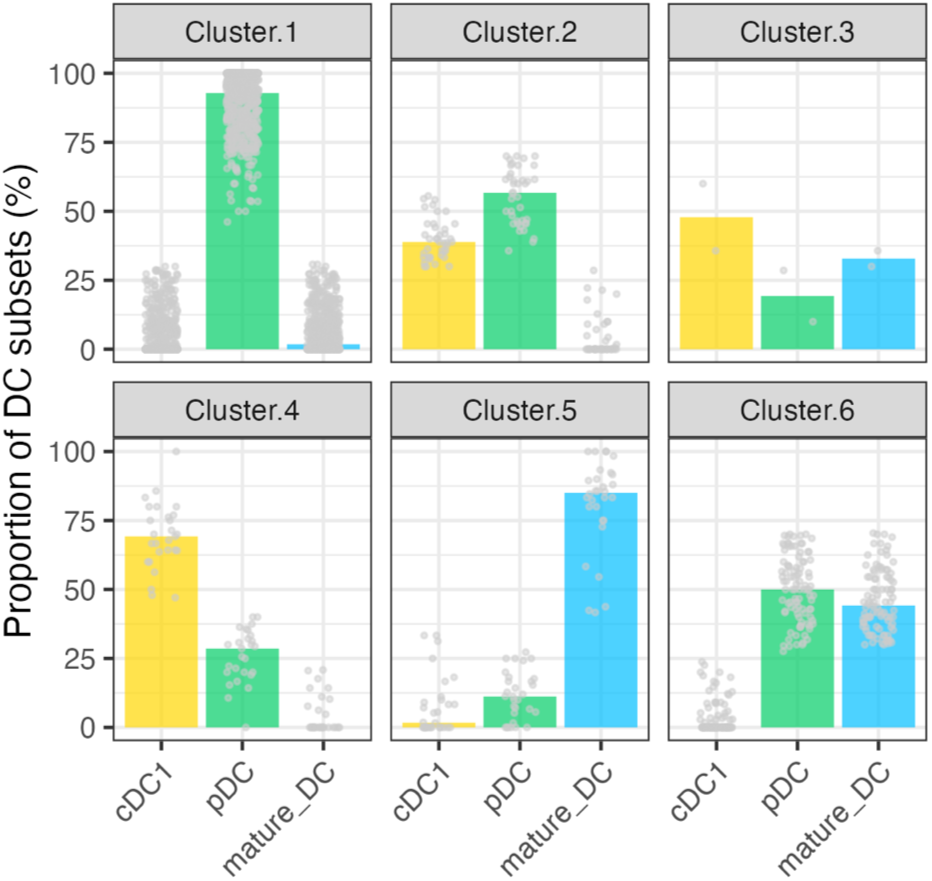
Composition of DC aggregates.

